# ERM proteins: The missing actin linkers in clathrin-mediated endocytosis

**DOI:** 10.1101/307272

**Authors:** Audun Sverre Kvalvaag, Kay Oliver Schink, Andreas Brech, Kirsten Sandvig, Sascha Pust

## Abstract

Canonical clathrin-coated pits (CCPs) are nucleated by the coordinated arrival of clathrin triskelia and AP2 adaptor proteins at phosphatidylinositol 4,5-bisphosphate (PIP_2_) enriched plasma membrane domains (1, 2). Subsequent propagation of the clathrin lattice helps to deform the membrane into a sharply curved pit (3). A large proportion of the initiated pits fall apart as abortive endocytic events within about 20 s, possibly due to insufficient cargo capture (4). Successful clathrin-mediated endocytosis (CME) is concluded when a clathrin-coated vesicle is released into the cell by dynamin-mediated fission of the CCP membrane neck (5, 6). A vast array of accessory proteins important for successful CME has been identified. Among these is actin, which has been shown to be required for the maturation and internalization of a subset of CCPs in human cells (7, 8). Actin dependency during CME correlates with elevated membrane tension, and CCP maturation requires actin polymerization during mitosis, in microvilli at the apical surface of polarized cells and in cells upon global mechanical stretching or hypotonic treatment (9–11). The Arp2/3 complex is thought to trigger an acute actin burst coinciding with CCP internalization, but how actin is recruited to CCPs in the first place is not known (12, 13). Here we show that the ERM (ezrin, radixin, moesin) protein family of membrane-actin linkers associates with CCPs, and that functional perturbation of ERM proteins impedes CCP maturation and reduces the rate of transferrin uptake. By total internal reflection fluorescence (TIRF) microscopy and unbiased object detection and tracking, we show that ezrin localizes to nascent CCPs and that these pits subsequently recruit actin. Based on these data, we propose a model in which activated ERM proteins recruit the initial actin filament during CME.

---

ERM proteins exist in the cytosol in a dormant, closed conformation in which the actin-binding C-terminal domain associates with a conserved N-terminal FERM (four-point-one, ERM) protein interaction domain. This conformation renders the proteins inactive as most protein binding sites are masked (14, 15). ERM proteins are activated upon interacting with PIP_2_ at the plasma membrane, followed by phosphorylation of a conserved threonine residue (Thr567, Thr564 and Thr558 for ezrin, radixin and moesin respectively) (16–20). This two-step activation mechanism induces a conformational change by which the proteins adopt an extended open conformation which exposes their protein binding sites. Through their FERM domain, activated ERM proteins can bind to transmembrane receptors, such as epidermal growth factor receptor (21) and alpha-1 adrenergic receptor (22), and membrane associated proteins, such as EBP50 (23). ERM proteins bind actin at their C-terminal domain. Thus, these proteins function as membrane-actin linkers involved in a wide range of functions, including regulation of cell morphology, cell migration and adhesion (24, 25). Moesin knockdown has been shown to inhibit clathrin dependent internalization of S1PR1 in T cells (26), but the role of ERM proteins in CME is not known.

Here we used genome-edited SUM159 breast cancer cells expressing endogenous AP2-eGFP to analyse native CME in real time by TIRF microscopy. AP2 is an integral component of CCPs and it is exclusively involved in CME (1, 27). Fluorescently tagged AP2 thus serves as an optimal marker for CME. We first used GFP-Trap^®^ to pull down AP2 and probed for the presence of ERM proteins (Figure 1a). All three ERM proteins could be detected in the precipitate and the level of co-immunoprecipitated proteins appeared to be decreased upon functional perturbation of ERM proteins by the small molecule inhibitor NSC668394 (described in (28), hereby termed ERM inhibitor (ERMi)). We proceeded by directly examining whether AP2 and ezrin colocalize at the plasma membrane by applying a fluorescence based proximity ligation assay (PLA), in which a fluorescent signal is emitted when two immunolabeled proteins are located within 40nm of each other (29). As shown in Figure 1b, ezrin and AP2 could be detected in the same membrane domains, and the number of detections was reduced after treatment with the ERM inhibitor. Next, we investigated the functional role of ERM proteins in CME by quantifying the rate of transferrin uptake and observed a decrease of about 80% after ERMi treatment (Figure 1c).

**Figure 1:**
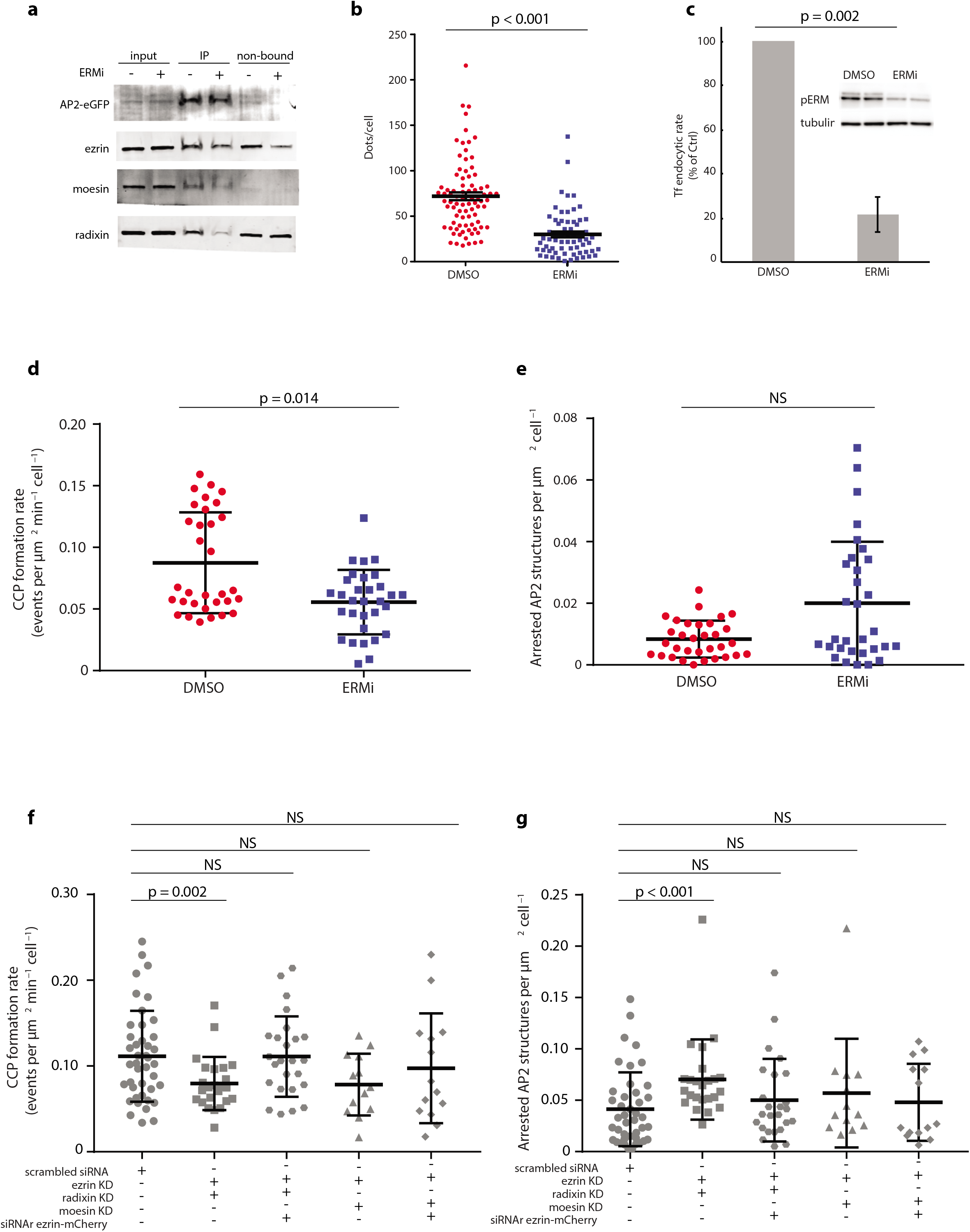
ERM proteins colocalize with CCPs and are involved in CME. **(a)** Representative western blot showing co-immunoprecipitation of AP2-eGFP and ERM proteins performed by GFP-Trap^®^ followed by Western blot analysis with corresponding antibodies. **(b)** Quantification of the association between AP2 and ezrin by Proximity Ligation Assay (PLA). SUMD8 cells were treated with DMSO or ERMi and ezrin-AP2 proximity signals were visualized by confocal microscopy and quantified with ImageJ. Each dot in the plots represents the number of detected proximity signals per cell. **(c)** Bar chart of cells treated either with DMSO or ERMi for 2 h, followed by 5 min incubation with ^125^I-transferrin (mean +/-SEM, n = 4). The insert shows the level of phosphorylated ERM proteins (pERM) in DMSO and ERMi treated cells **(d)** Scatter plots of the number of bona fide CCPs formed per μm^2^ per min per cell captured from initiation to internalization during 5 min movies at 0.5 fps in cells treated with DMSO or ERMi for 2 h (mean +/-SEM, n = 32 DMSO and n = 31 ERMi). **(e)** Scatter plots of the number of arrested AP2 structures per μm^2^ per cell with lifetimes exceeding the duration of the movies (> 5 min) (mean +/-SEM). **(f)** Scatter plots of the number of CCPs formed in cells treated with scrambled siRNA compared to cells transfected with a combination of siRNAs targeting either ezrin and radixin (EzRa) or ezrin and moesin (EzMo), and siRNA treated cells expressing siRNA resistant ezrin-mCherry in addition (rEzRa/rEzMo, mean +/-SEM, n_cells_ = 46 scr, n_cells_ = 28 EzRa, n_cells_ = 21 rEzRa, n_cells_ = 25 EzMo and n_cells_ = 23 rEzMo). **(g)** Scatter plots of the same cells as in **(f)**, but here showing the number of arrested AP2 structures per μm^2^ per cell (mean +/-SEM).

To further investigate the functional relationship between ERM proteins and AP2, we quantified the effect of ERM inhibition on the number of CCPs formed at the basal surface of cells (i.e. the CCP formation rate). To this end, we captured movies (5 min, 0.5 frames per second (fps)) of the SUMD8 cells by TIRF microscopy and analysed the movies by applying an automated CME analysis algorithm enabling unbiased CCP detection and tracking, as published previously (30). By comparing the rate of bona fide CCP formation per μm^2^ in cells treated either with the ERM inhibitor or DMSO, we observed a mean reduction of about 40% (Figure 1d). We also quantified the number of arrested AP2 structures, with lifetimes exceeding 5 min, which increased in about 50% of the ERMi treated cells. However, the mean difference across the population was not significant (p = 0.0587). To verify that these effects were in fact induced by ERM protein inactivation, we treated cells with siRNAs targeting combinations of the different ERM protein family members and quantified the CCP formation rate. By comparing control cells treated with non-targeting siRNA to double knockdown cells treated with siRNA oligos targeting ezrin and radixin, we observed a ~30% reduction (Figure 1f). We observed a similar tendency after ezrin and moesin double knockdown, although this effect was considered as not to be significant (p = 0.06). We could also observe a significant increase in the number of arrested structures following ezrin/radixin double knockdown (Figure 1g). These effects could be rescued with a siRNA-resistant ezrin construct (Figure 1f-g). We also performed triple knockdown of all ERM proteins, but the treatment strongly affected cell proliferation and viability and was not analysed further (data not shown).

Our results demonstrate that ERM proteins are involved in CCP formation and/or maturation. Based on the well-established role of the ERM proteins as PIP_2_-interacting membrane-actin linkers, we proceeded by investigating the functional relationship between ERM proteins and actin during CME. We first analysed to which extent actin polymerization is required for CCP maturation in these cells by treating them with 1 μM of the F-actin stabilizing drug Jasplakinolide (Jasp). We then observed a 60% reduction in the CCP formation rate, accompanied by a 6-fold increase in the number of arrested AP2 structures (Figure 2a-b). These data indicate that dynamic actin is important for the maturation of a subset of the CCPs in these cells.

**Figure 2:**
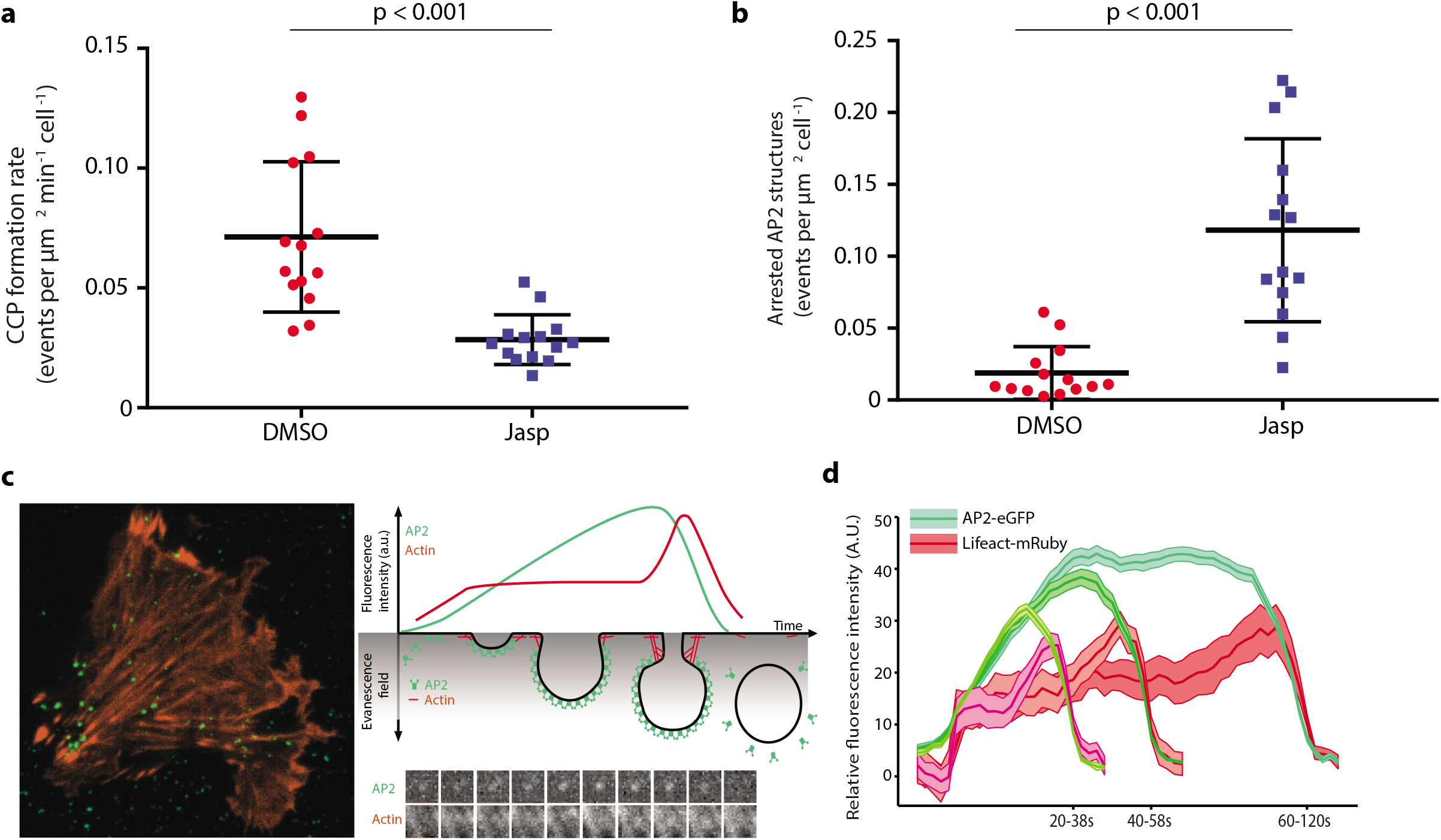
Actin involvement in CME. **(a)** Scatter plots of the number of CCPs formed in cells treated with DMSO compared to cells treated with 1μM Jasplakinolide (Jasp) for 15-45 min (mean +/-SEM, n_cells_ = 14 per treatment). **(b)** Scatter plots of the same cells as in **(a)**, but here showing the number of arrested AP2 structures per μm^2^ per cell (mean +/-SEM). **(c)** Left panel: A TIRF micrograph of a SUMD8 cell expressing AP2-eGFP and Lifeact-mRuby. Upper right panel: A schematic illustration of TIRF imaging of the temporal actin and AP2 recruitment during formation of a CCP, and the corresponding fluorescence intensity traces. Lower right panel: A representative frame-by-frame display of AP2-eGFP and Lifeact-mRuby recruitment. **(d)** Temporal fluorescence intensity traces distributed in three lifetime cohorts of AP2-eGFP and Lifeact-mRuby from CCPs in which the Lifeact intensity exceeded the local background signal for > 6 consecutive seconds (mean +/-SEM, n_cells_ = 12).

Next, we analysed the temporal dynamics of actin recruitment to CCPs by capturing 5 min movies at 2 fps by two channel TIRF microscopy of SUMD8 cells expressing the fluorescently tagged actin-binding peptide Lifeact-mRuby and AP2-eGFP (Figure 2c). When we subsequently applied a minimal fluorescence intensity threshold corresponding to a Lifeact intensity greater than the local background for 6 consecutive seconds, we detected actin recruitment in about 60% of the CCPs, which is in the same range as reported previously (8). We also detected actin recruitment in two distinct phases, one early phase during which a plateau is reached within the first third of the CCP lifecycle, and one late phase with a massive actin polymerization coinciding with CCP internalization (Figure 2d). These data are consistent with a model where primary actin filaments are recruited during the initial phase of CCPs maturation to act as “mother” filaments, which are required as platforms for subsequent Arp2/3-triggered actin polymerization during CCP internalization.

To simultaneously visualize the spatiotemporal dynamics of AP2 and members of the ERM protein family, we created double genome-edited cells expressing endogenous levels of AP2-eGFP and ezrin-mScarlet (hereby termed SUMD8e, described in Supplementary Fig. 1). By capturing 3 min movies at 1/3 fps of these cells by two-color TIRF microscopy, we could follow individual CCPs and correlate the temporal ezrin-mScarlet fluorescent intensity to AP2-eGFP. We subsequently filtered bona fide CCP tracks by applying a minimal ezrin-mScarlet intensity threshold over the local background for > 6 consecutive seconds, as before. We then observed that the bulk ezrin recruitment occurred early during CCP formation, with a smaller secondary burst coinciding with the CCP internalization phase (Figure 3a). Figure 3b shows the temporal Lifeact recruitment from Figure 2d, plotted in individual lifetime cohorts. To compare these data with the temporal recruitment of other known actin binding proteins involved in CME, such as Hip1R, dynamin2 and Arp3 (from the Arp2/3 complex), we imaged cells transiently expressing low levels of fluorescently tagged versions of these proteins in the SUMD8 cells and analysed their temporal intensity profiles relative to AP2 as before. In contrast to ezrin, Hip1R, dynamin2 and Arp3 showed a later peak of the fluorescence intensity, contemporary with the AP2 peak around the internalization phase (Figure 3c-e).

**Figure 3:**
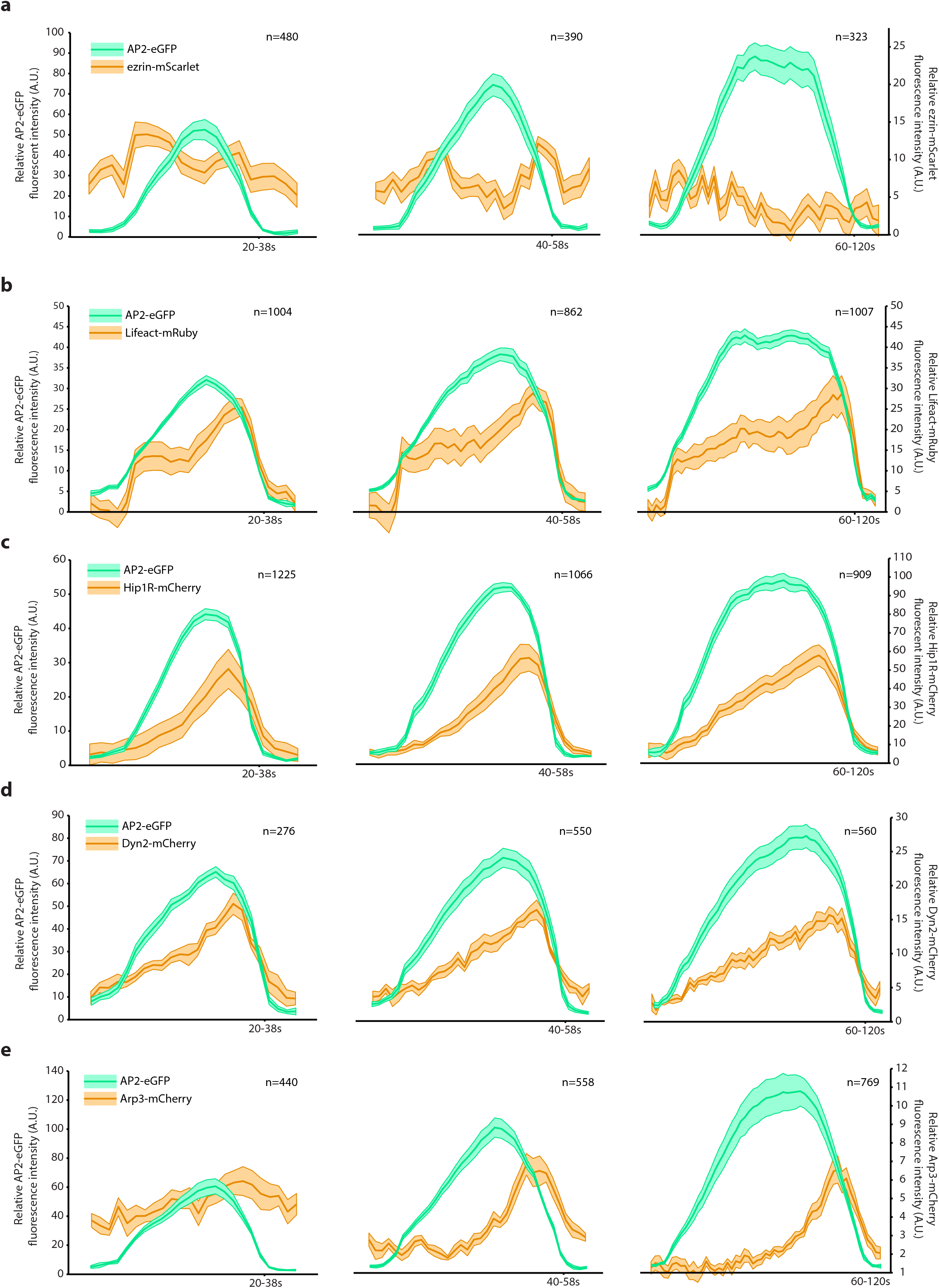
Endocytic accessory protein (EAP) recruitment during CME. Endocytic events are separated into three lifetime cohorts (20-38 s, 40-58 s and 60-120 s), and the number of analysed tracks per lifetime cohort per EAP is indicated in the top right corner of each plot. **(a)** Temporal fluorescence intensity traces of AP2-eGFP and ezrin-mScarlet from CCPs in which the ezrin-mScarlet intensity exceeded the minimal ezrin-mScarlet intensity threshold for > 6 consecutive seconds (mean +/-SEM, n_cells_ = 10). **(b)** Separated lifetime cohorts of the fluorescence intensity traces of AP2-eGFP and Lifeact-mRuby shown in Figure 2d. (**c-e**) Temporal fluorescence intensity traces (mean +/-SEM) of AP2-eGFP with **(c)** Hip1R-mCherry (n_cells_ = 5), **(d)** dynamin2-mCherry (n_cells_ = 6) and **(e)** Arp3-mCherry (n_cells_ = 6).

Next, we transiently transfected the SUMD8e cells with a SNAP-tagged silicone rhodamine 647 (SNAP-Cell^®^ 647-SiR) Lifeact (Lifeact-SiR-647) construct to simultaneously capture the temporal fluorescence intensity profiles of AP2-eGFP, ezrin-mScarlet and actin (Lifeact-SiR-647) by three-channel TIRF microscopy. We captured 3 min movies with 1/3 fps framerates and applied a minimal fluorescence intensity threshold for Lifeact-SiR-647 to the acquired CCPs for a minimum of 9 consecutive seconds. We then observed a similar intensity profile for ezrin as when we applied an ezrin-mScarlet threshold, in which the peak of ezrin recruitment correlated temporally with the initiation of the actin plateau (Figure 4a). To analyse to which extent ezrin recruitment correlated with actin recruitment, we analyzed CCPs that recruited Lifeact-SiR-647 according to whether they also reached an ezrin-mScarlet minimal intensity threshold (Figure 4b-c). We then observed that actin recruitment correlated with ezrin recruitment in ~70% of the CCPs. When we compared the ezrin-mScarlet intensity of the pits exceeding the Lifeact-SiR-647 threshold to the pits that failed to reach the threshold, and we observed only nominal ezrin recruitment across all lifetime cohorts among the pits not recruiting actin (Figure 4d). Taken together, these data demonstrate a direct correlation between ezrin and actin recruitment during CME. To further investigate the structural organization of actin, ezrin and AP2, we performed structured illumination microscopy (SIM) of SUMD8 cells with fluorescently labelled phosphorylated ERM proteins (pERM) and actin, in addition to AP2-eGFP. Figure 4e and the corresponding Supplementary Movie S1 show an example of a CCP associated with pERM and actin.

**Figure 4:**
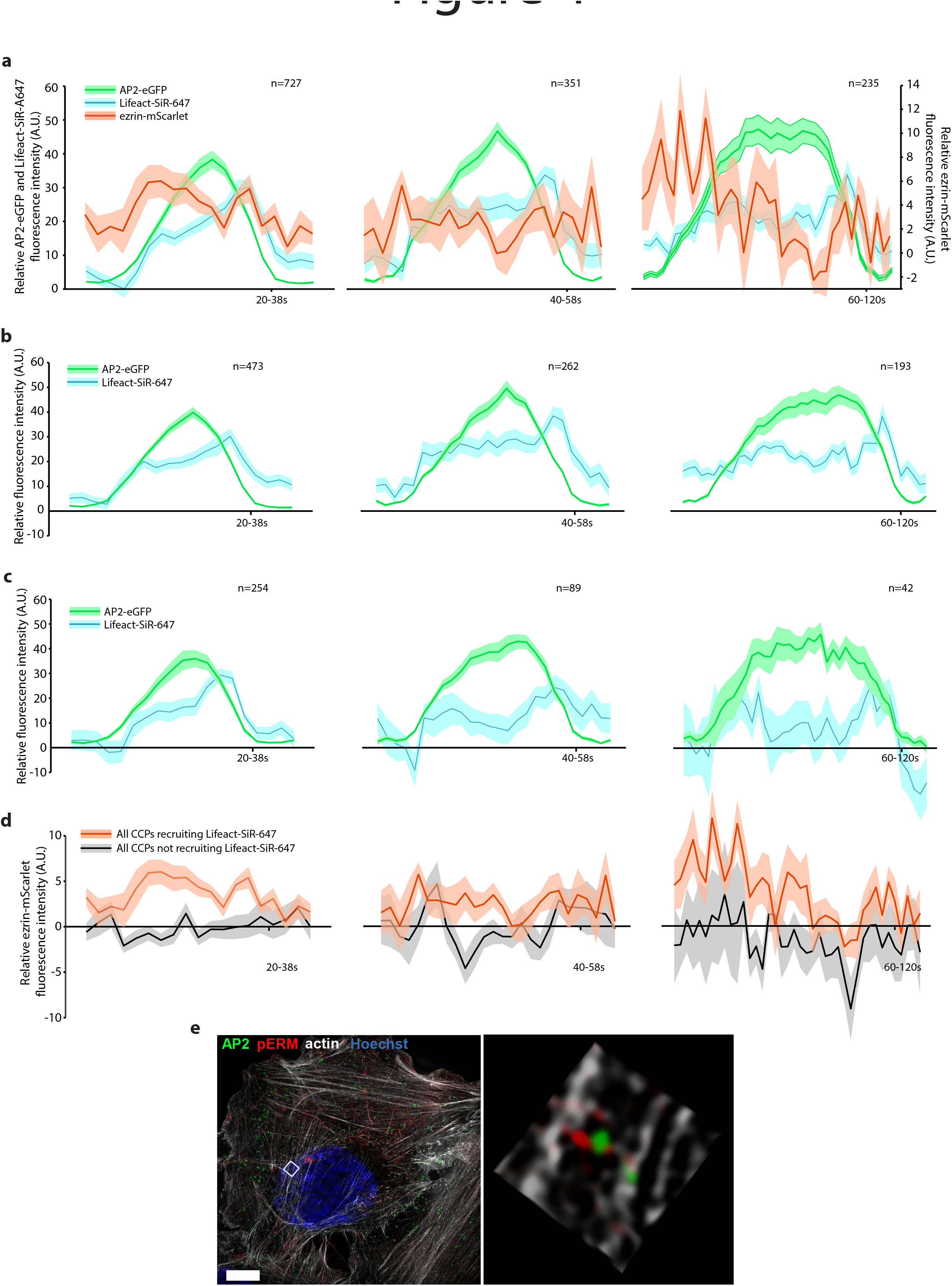
Ezrin and actin recruitment during CME. Endocytic events are separated into three lifetime cohorts (20-38 s, 40-58 s and 60-120 s), with the number of analysed tracks indicated in the top right corner of each plot. **(a)** Temporal fluorescence intensity traces of AP2-eGFP, ezrin-mScarlet and Lifeact-SiR-647 from CCPs in which the Lifeact intensity exceeded the local background intensity for > 9 consecutive seconds (mean +/-SEM). **(b)** The temporal fluorescence intensity traces of AP2-eGFP and Lifeact-SiR-647 from CCPs filtered in which the ezrin-mScarlet intensity exceeded the minimum intensity threshold for > 9 consecutive seconds (mean +/-SEM) **(c)** The temporal fluorescence intensity traces of AP2-eGFP and Lifeact-SiR-647 from CCPs in which the ezrin-mScarlet intensity failed to reach the threshold from **(b)**. **(d)** Temporal ezrin-mScarlet fluorescence intensity traces from CCPs in which actin was detected compared to the ezrin-mScarlet intensity in CCPs with no actin recruitment (mean +/-SEM). **(e)** Left panel: Micrograph showing the maximum intensity projection of a Z-stack acquired by 4 color super-resolution 3D-SIM with 125 nm increments of a SUMD8 cell stained for pERM and actin, and with the nuclear marker Hoechst (scale bar = 5 μm). Right panel: An enlarged version of the highlighted region in the micrograph.

Actin has been shown to be required for CME under certain circumstances, such as in membranes with high tension (9–11) and during steric hindrance of clathrin coat assembly (31, 32). Here we show that activated ERM proteins associate with CCPs, and that they are required for efficient CME. Ezrin associates with CCPs during their initiation, and ezrin recruitment reaches a maximum within the first third of CCP maturation. This correlates temporally with initial actin recruitment. In contrast with the temporal ezrin recruitment, other actin binding proteins involved in CME, such as dynamin and Hip1R, accumulate during CCP maturation and peak around CCP internalization. These temporal recruitment profiles do not preclude these proteins from being involved in actin recruitment, but they might indicate a mechanism by which these proteins prime CCPs for the acute Arp2/3 driven actin burst occurring during CCP internalization. Finally, we show direct evidence for a strong correlation between ezrin recruitment and actin recruitment to CCPs in minimally perturbed cells. Based on these data, we propose a model in which ERM proteins associate with nascent CCPs to recruit primary actin filaments to provide the “mother” filaments needed for subsequent Arp2/3 driven actin polymerization during CCP internalization.

## Materials and methods

### Cell culture

Gene-edited SUM159PT human breast cancer cells expressing either AP2 σ2-eGFP alone (here termed SUMD8 described in (33) as SUM-AP-2.1), or both AP2 σ2-eGFP and ezrin-mScarlet (SUMD8e) were cultured in Ham’s F-12 culture medium supplemented with HEPES (10mM), Insulin (5μg/ml), Hydrocortisone (1μg/ml) and 5% fetal bovine serum (FBS) at 37°C and 5% CO_2_. Sub-confluent cells were plated in appropriate culture vessels one day prior to experiments. SUMD8 cells were used unless otherwise stated.

### Antibodies and other reagents

Antibodies used for immunoblotting: mouse anti-ezrin (E8897, Sigma Aldrich), rabbit anti-moesin (3146, Cell Signaling), rabbit anti-radixin (HPA000763, Sigma Aldrich), rat anti-GFP (Chromotek) and mouse anti-GAPDH (Ab-9484, Abcam). For microscopic analysis rabbit anti-pERM (48G2, Cell Signaling) and Phalloidin-647 was from Molecular Probes. Jasplakinolide was purchased from Abcam, *n*-octylglucopyranoside were purchased from Sigma-Aldrich, and the small molecule inhibitor NSC668394 was purchased from Calbiochem.

### Transfection of siRNA oligos and plasmids

Cells were plated 24 h prior to transfection. They were then transiently transfected with 25 nM siRNA oligos (Dharmacon) using DharmaFECT 1 transfection reagent (Dharmacon) according to the manufacturer’s protocol. Knockdown of ezrin and moesin was performed with oligos described in (34) and for radixin knockdown oligos with the target sequence CUACAUGGCUUAAACUAAA was used. A non-targeting siRNA oligo was included as a control in all siRNA knockdown experiments. Cells were then grown for 72-84 h before experimental procedures were carried out.

For rescue experiments, cells were grown for 24 h after siRNA transfection. Then they were transfected with 2 μg of plasmid carrying siRNA resistant ezrin (described in (34)) per 8 μl FuGENE HD transfection reagent (Roche) according to the manufacturer’s protocol. Control cells were transfected with an empty mammalian expression vector. The cells were then grown for additional 24-36 h prior to experimental procedures. All other plasmid transfections were carried out 24 h after the cells were plated, followed by 48 h plasmid incubation. Transfected cells were washed and incubated in fresh full SUM159 culture medium > 4 h prior to imaging.

### Cell lysis and Western Blotting

Cells were washed in cold PBS and lysed in lysis buffer containing 0.1 M NaCl, 10 mM Na_2_HPO_4_, 1 mM EDTA, 1% Triton X-100, 60 mM n-octylglucopyranoside, pH 7.4 and supplemented with complete protease inhibitors and PhosSTOP (Roche Diagnostics). The lysate was cleared by centrifugation and proteins were separated by SDS-PAGE on a 4-20% gradient gel (Protean-TGX, Bio-Rad) and blotted onto a PVDF membrane (Bio-Rad). The membrane was blocked in 5% w/v BSA (Sigma) in PBS with 0.1% Tween-20 (PBS-T) before incubating with the primary antibody overnight at 4 °C. After washing with PBST, IRDye infrared linked secondary antibodies (LI-COR Biosciences) were used according to the manufacturer's manual The bands were detected by Odyssey Infrared Imaging System (I I_-_COR Biosciences) and quantified by using FIJI/ImageJ. Levels of GAPDH were used as internal loading controls and data were further correlated to those of the control treatments.

### Immunoprecipitation

Co-immunoprecipitation experiments of AP2-GFP were performed by using GFP-Trap^®^ as described in the manufacturer’ s instructions. SUMD8 cells were treated with the ERM inhibitor (2h, 10 μM), lysed [10 mM Tris pH 7.5, 150 mM sodium chloride, 0.5 mM EDTA, 2% NP40, protease inhibitor cocktail (complete, Roche)], pelleted and the supernatant incubated with GFP-Trap agarose beads (Chromotek) as described in the manufacturer’ s instructions. Protein samples of the total lysate (input), IP fractions and the non-IP fractions (non-bound) were separated by SDS-PAGE and blots were probed with corresponding antibodies.

### Live cell TIRF imaging

Cells were seeded in sterile 35 mm glass bottom culture dishes (MatTek Corporation) at a density yielding about 70% confluency at the time of imaging, and maintained in full SUM159 culture medium. Just prior to imaging, the culture medium was exchanged with pre-warmed (37°C) α-MEM without phenol red (Gibco) supplemented with 5% FBS and 10 mM Hepes buffer (Sigma-Aldrich). TIRF movies were acquired using a Deltavision OMX V4 system (Applied Precision, GE Healthcare) with a ring TIRF design (licensed to GE Healthcare from Yale University) rotating the laser beam at the back focal plane of the objective lens. This generates the TIRF evanescence field almost simultaneously from multiple points, resulting in highly uniform TIRF illumination. The microscope was equipped with a 60x ApoN NA 1.49 objective (Olympus), three cooled sCMOS cameras (PCO) and 488nm (100mW OPSL), 568 nm and 642 nm (110 mW diode) lasers. The stage and objective were temperature controlled at 37°C and the sample holder was enclosed by a humidified 5% C0_2_ incubation chamber. The system was controlled using the OMX Acquisition control software running under a Windows 7 operating system. Image registration was performed using the SoftWoRx software package (Applied Precision, GE Healthcare) The final pixel size was 80 nm.

### Data analysis of TIRF movies

Post-processing of raw data was carried out in the Matlab environment (Mathworks), essentially as described by Aguet *et al.* in (30). However, Aguet *et al.* determines association between fluorescently tagged endocytic accessory proteins (EAPs) and CCPs by statistical testing of the EAP fluorescent signal within the CCP mask compared to the signal excluded from the mask. In our hands, this approach becomes arbitrary when analyzing EAPs such as actin and ezrin that are highly expressed in non-CCP membrane domains. We therefore applied a fixed threshold over a given time interval to determine association for all EAPs examined in this study. However, it should be noted that we obtained similar temporal fluorescence intensity profiles by applying statistical testing for all EAPs except actin.

### Duolink - in situ proximity ligation assay

The Duolink© proximity ligation (PLA) technology (Sigma-Aldrich) was used to analyze and quantify protein-protein interactions. The cells were washed in PBS and fixed in 4% PFA for 15 min, permeabilized with 0.1% Triton X-100 in PBS for 2 min. Primary antibodies raised against GFP and ezrin, were used and the PLA reactions were performed as described in the manufacturer’s instructions. The Samples were mounted with Duolink Mounting Medium ProLong Diamond with DAPI (Invitrogen) and images were acquired by a LSM-780 confocal microscope (Carl Zeiss) and analyzed with FIJI software (35). For the negative controls, only one antibody or antibodies targeting non-interacting proteins were used. Mitotic and apoptotic cells were excluded from the analysis.

### 3D-SIM

For 3D-SIM (structured illumination microscopy) cells were fixed 8% PFA for 15 min (36) and permeabilized with 0.1% Triton X-100 in PBS for 5 min. For protein staining, a specific primary antibody against pERM (#2217) and Alexa647-conjugated phalloidin was used. Coverslips were mounted with ProLong Glass (Invitrogen). 3D-SIM imaging was performed on a Deltavision OMX V4 system (Applied Precision) equipped with an Olympus 60x numerical aperture (NA) 1.42 objective, cooled sCMOS cameras and 405, 488, 568 and 642 nm diode lasers. Z-stacks covering the whole cell were recorded with a Z-spacing of 125 nm. A total of 15 raw images (five phases, three rotations) per plane were collected and reconstructed by using SOFTWORX software (Applied Precision) and processed in FIJI, ImageJ or icy software.

### CRISPR-mediated generation of SUMD8e cells

The genome-edited SUMD8e cell line was generated by fusion of ezrin to the mScarlet flourescent tag, based on site directed introduction of CRISPR/Cas9n-targeted DNA breaks and template assisted homology driven repair. A CRISPR/Cas9 nickase strategy was used to fuse mScarlet to the C-terminus of ezrin, by targeting the end of exon 13 of the ezrin gene, inserting a 13 amino acid linker (SGGSGGGSGPVAT) and mScarlet. In the donor plasmid, this inserted sequence was flanked by 1k base pair homology arms for HDR. The homology arms, linker region and mScarlet was synthesized (GeneArt^®^, Invitrogen) and cloned into pMS-RQ vector (Invitrogen). The sgRNA was designed using the Benchling online tool with different oligos (Table S1) and prepared as described in (37) and cloned into the px459 vector. Cells were selected by using 1-10 μg/ml puromycin for 3 days, until all non-modified cells died. Positive cells were further expanded to confluency in culture flasks and FACS single-cell sorted for mScarlet levels. Single cell clones were expanded and characterized.

### Wound healing assay

Cells were seeded on Glass Bottom Dishes (MatTek Corporation) and grown until confluency. Cells were then scratched with a 10 μl tip and subsequently observed with a BioStation IM Live Cell Recorder (Nikon Instruments Inc.). Image acquisition was done every 5 min for 7 h in total. Images were analysed with ImageJ software with Manual Tracking and Chemotaxis and Migration Tool (ibidi GmbH) plugins. Directionality is calculated by dividing the Euclidean distance by the accumulated distance of the cell trajectories.

### Transferrin (Tf) uptake

Tf was labelled with I^125^ using Pierce Iodination Tubes (Thermo Fisher) according to the manufacturer’s protocol to a specific activity of about 25000 cpm/ng. Cells were seeded in 24 well plates and washed once with Hepes buffered medium. The cells were then treated with 10 μM ERMi in 200 *μ* L HEPES-buffered medium for 2 h at 37°C. Cells were then incubated with 40 ng ^125^l-Tf per well in a total volume of 200 *μ* L for 6 min at 37°C. The cells were then washed 3 times with ice cold PBS and incubated on ice in the presence of 2 mg/ml_ protease Pronase E (Sigma-Aldrich) in 300 *μ* L HEPES-buffered medium for 1 h, which enables us to separate internalized and membrane-bound Tf. The cells and the supernatant were subsequently transferred to Eppendorf tubes and centrifuged at 14 000 rpm for 2 min. The radioactivity in the supernatant (cell-surface associated Tf) and pellet (internalized Tf) was quantified using a LKB Wallac 1261 Multigamma counter (LKB Instruments) and endocytosis was expressed as the amount of internalized Tf divided by the total amount of cell-associated Tf.

### Statistics

Statistical significance of TIRF data was tested by the Wilcoxon rank sum test. Student's t-test or Mann-Whitney rank sum test was used to calculate the P-value for all other experiments. A P-value of 0.05 or less was considered to be statistically significant. A minimum of 3 independent experiments were performed and quantification of the data is given as mean ± SD.

## Acknowledgements

We wish to thank Anne-Grethe Myrann for technical assistance and Anne Engen and her team for excellent assistance with cell culturing. The gene-edited SUMD8 cells were a kind gift from Prof. Tom Kirchhausen at Harvard Medical School/PCMM, Boston, MA, USA. This work was supported by the Norwegian Cancer Society, the South-Eastern Norway Regional Health Authority and the Research Council of Norway through its Centres of Excellence scheme, Project Number 17957.

